# Recursive Entropic Time: A Neural Framework for the Informational Construction of Subjective Duration

**DOI:** 10.1101/2025.07.03.663003

**Authors:** Khaled Bouzaiene

## Abstract

The conception of time as a universal, independent parameter is a foundational axiom of physical models, yet it fails to account for the subjective nature of temporal perception and leads to theoretical inconsistencies in complex systems. This paper introduces and provides multifaceted empirical support for the Recursive Entropic Time (RET) framework, a theory positing that subjective time is not a fixed background but an emergent property actively constructed by neural systems engaged in interpretive, associative processing. We hypothesize that the brain employs a functionally segregated system for temporal processing: primary sensory cortices negotiate objective, clock-based time (*t_lab_*), while higher-order associative cortices construct subjective time (*t_RET_*) via a mechanism where the rate of temporal flow is inversely modulated by moment-to-moment informational load.

We tested this theory through a comprehensive, two-part investigation. Part I utilized a public EEG-fMRI dataset of subjects under the influence of DMT to test the anatomical specificity of RET. This analysis revealed a crucial discovery: the RET model’s efficacy, quantified by the improvement in correlation between neural dynamics and informational load, was significantly greater in associative regions (e.g., Lingual Gyrus) compared to primary sensory regions (V1) (p = 0.0068). This finding refuted a global model of RET and pointed toward a more nuanced, region-specific hypothesis. Part II was designed to probe the unique, non-linear dynamics predicted by this refined hypothesis. We conducted a mechanistic analysis on a public EEG dataset of a temporal reproduction task (Hassall et al., 2020), identifying two behaviorally identical trials (reproduced duration *≈* 0.78s). We demonstrated that their underlying informational load profiles were demonstrably different, a finding inconsistent with simpler linear scaling models. Crucially, the formal RET integral correctly predicted the behavioral outcome, yielding nearly identical total internal durations.

Together, these results provide convergent evidence for RET, presenting it as a falsifiable and mechanistic account of how the brain constructs subjective time. We argue that time, as we experience it, is not a perception of an external property but an intrinsic construction of higher-order cognition. This work establishes a computationally tractable and falsifiable method for quantifying subjective time from first principles, with implications for creating dynamic “brain-time” atlases and understanding cognitive pathologies.

## 1 Introduction

### 1.1 The Two Clocks: The Discrepancy Between Physical and Psychological Time

The nature of time remains one of the most profound and unresolved questions at the intersection of physics, neuroscience, and philosophy. In the paradigms of classical mechanics, computation, and even much of quantum mechanics, time is treated as a smooth, immutable, and universally shared background parameter—a “conveyor belt” upon which events unfold [1]. This idealized, absolute time is a powerful and necessary construct for modeling the objective, physical world. Its success is undeniable, forming the basis for technologies that require picosecond precision. However, its rigid structure creates a deep chasm between our physical models and the lived reality of psychological time, which is notoriously fluid, subjective, and state-dependent [2, 3]. The subjective feeling of time “flying” during moments of engagement or “dragging” during periods of boredom or fear is a universal human experience that a simple, ticking clock fails to explain. This duality presents a central challenge: how does the seemingly objective, clock-like progression of the physical world give rise to the malleable, story-like progression of conscious experience?

### 1.2 Existing Models in Neuroscience: From Pacemakers to Scaling Laws

The search for the neurobiological basis of time perception has evolved significantly. Early “pacemaker-accumulator” models posited a centralized internal clock that emits pulses counted by a separate module [4]. However, the search for a single, dedicated “pacemaker” in the brain has been largely unsuccessful. This has led to the current consensus that timing is a distributed, intrinsic property of neural network dynamics [6, 5]. In this view, the state-dependent firing patterns and synaptic plasticity of neural populations are themselves the mechanism by which the brain tracks time.

A significant recent advancement in this area is the “Temporal Scaling” hypothesis, elegantly demonstrated by Hassall et al. (2020) [7]. By analyzing EEG data from a time estimation task, they showed that the trial-to-trial variability in a subject’s behavior could be explained by linearly stretching or compressing the underlying neural activity (specifically, event-related potentials). This suggests the brain can globally “speed up” or “slow down” a neural process for an entire event, as if adjusting a playback speed. This is a powerful descriptive model that successfully accounts for behavioral variability. However, it treats the scaling factor k as a single, static parameter for the whole event, leaving unanswered the deeper, mechanistic questions: What biophysical process does k represent? What causes it to change from one trial to the next? And is it possible for the speed of internal time to change within a single event?

### 1.3 The Recursive Entropic Time (RET) Hypothesis: A Mechanistic Proposal

This paper introduces a novel theoretical framework, Recursive Entropic Time (RET), which posits a more granular and mechanistic solution. RET proposes that subjective time is not merely scaled globally but is actively and locally constructed from moment to moment. The central thesis is that the rate of progression of a system’s internal, subjective time is inversely proportional to its instantaneous “informational load.”

Informational load, *L*(*t*), is formally defined as the magnitude of change in the system’s informational state, S(t), a proxy for “surprise” or the computational effort required to update an internal model of the world, consistent with predictive coding frameworks [8]. The mathematical formulation is:

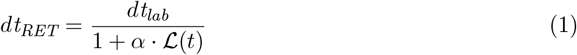

When the brain encounters a predictable, low-complexity informational environment, *L*(*t*) is low, and the internal clock runs quickly (*dt_RET_* ≈ *dt_lab_*). When it encounters a novel, complex, or rapidly changing environment, ℒ(*t*) is high, and the clock slows down (*dt_RET_* < *dt_lab_*), dedicating more “internal moments” to resolve the complexity. Time, in this view, is the very process of a system metabolizing information.

This framework allows us to make a unique, falsifiable prediction that distinguishes it from global scaling models. While a scaling model would predict that two behaviorally identical events must have identically scaled neural dynamics, RET predicts that two events with the same final duration could arise from vastly different internal temporal journeys. One event might have high informational load at the beginning and low load at the end; another could have the reverse profile. As long as the integral of their moment-to-moment temporal progression sums to the same total, the behavioral outcome will be the same. This non-obvious “equifinality” is the signature of a truly dynamic, emergent temporal process.

### 1.4 An Iterative Investigative Approach: From Anatomical Specificity to Mechanism

The validation of the RET framework, as detailed in this paper, was not a linear path but a process of iterative discovery. We designed a two-part investigation where the results of the first experiment directly motivated the design of the second.

Experiment 1 began with a broad, exploratory question: Does the RET model apply globally to the brain, especially in a state of known temporal distortion? We analyzed a public EEG-fMRI dataset of subjects under the influence of DMT. The initial results represented a crucial “failure” of the global hypothesis, as the RET model’s efficacy was not uniform. However, this finding led to the discovery of a striking, region-specific pattern, prompting a refined hypothesis of a functional segregation of temporal processing.

Experiment 2 was designed specifically to test this new, more nuanced hypothesis and to probe the unique, non-linear mechanism of RET. We selected a different public EEG dataset from a temporal reproduction task, which allowed us to perform a definitive mechanistic analysis. By identifying two behaviorally identical trials, we could directly test the prediction that their internal informational dynamics would be different, a finding that would be inconsistent with simpler linear scaling models. This paper presents the full narrative of this scientific journey, demonstrating how the initial discovery of anatomical specificity led to a powerful, mechanistic validation of the RET framework.

## 2 Experiment 1: Discovering the Functional Segregation of Temporal Processing

### 2.1 Methods

#### Dataset and Code Availability

We utilized the publicly available simultaneous EEG-fMRI dataset from Timmermann et al. (2023) [11]. Data from 14 healthy participants who received an intravenous infusion of DMT were analyzed. This dataset is particularly suitable as psychedelic states are strongly associated with altered time perception and increased brain entropy [12]. All analysis code developed for this paper is available upon reasonable request.

#### Data Acquisition and Preprocessing

As detailed in the original publication [11], fMRI data were acquired on a 3T Siemens Verio scanner with a TR of 2000 ms. Standard preprocessing steps including slice-timing correction, realignment, normalization to MNI space, and spatial smoothing were applied. Simultaneous EEG data were recorded from a 64-channel system, and for our analysis, we used the pre-computed Lempel-Ziv complexity metric which was derived from the cleaned EEG signal.

#### Signal Definition

For each of the 14 subjects, two key time series were defined:

- **Informational State Signal:** This signal was derived from the Lempel-Ziv complexity data provided in the `RegressorsInterpscrubbedIRASA_Central.mat` file [13]. Lempel-Ziv complexity is a non-parametric measure of a signal’s compressibility and serves as a proxy for its entropy or information content. For each subject, the signal was averaged across its five provided components to create a single entropy vector.
- **Physical State Signal:** This signal was derived from the `BOLD_AAL` variable, representing the BOLD fMRI signal parcellated into 112 anatomical regions according to the Automated Anatomical Labeling (AAL) atlas.

#### Network and Regional Definition

To test for anatomical specificity, we conducted two levels of analysis. First, for a broad comparison, we defined three functional networks based on AAL atlas indices (0-based): Primary Visual Cortex (V1) [48, 49]; Lingual Gyrus [50]; and the Fronto-Parietal Network (FPN) [7, 8, 11, 12, 63, 64]. Second, to perform a more granular “deep dive” analysis within the visual system, we individually assessed all major regions of the visual cortex as defined by the AAL atlas: Calcarine [48], Cuneus [49], Lingual [50], Superior Occipital [51], Middle Occipital [52], Inferior Occipital [53], Fusiform [54], Superior Parietal [55], and Inferior Parietal [56]. For network-level analyses, BOLD signals were averaged across the constituent region indices.

#### Statistical Analysis

We compared two models of brain-information dynamics:

- **Lab Time Model:** This standard model calculates the absolute Pearson correlation between the raw informational state and the raw neural activity: |*r*|_Lab_ = |Pearson(Entropy(*t*), BOLD(*t*))|.
- **RET-Inspired Model:** This model tests the core tenet of RET by calculating the absolute Pearson correlation between the rates of change of both signals, approximated by the absolute first-order difference: |*r*|_RET_ = |Pearson(|*d*(Entropy)*/dt*|, |*d*(BOLD)*/dt*|)|.

The improvement in explanatory power was quantified as Δ|*r*| = |*r*|_RET_ − |*r*|_Lab_. To test our hypothesis of functional segregation, a two-sided paired t-test was used to compare the Δ|*r*| values between the Lingual Gyrus and V1 networks across all subjects, with a significance level of *p <* 0.05. All analyses were restricted to the first 5 minutes (150 fMRI timepoints) to capture the peak psychedelic effect.

#### 2.2 Results

#### 2.2.1 Broad Network Segregation

The primary analysis revealed a clear and statistically significant functional segregation. As shown in Figure 1a, the median improvement (Δ|*r*|) for the aggregated V1 network was strongly negative. In stark contrast, the median improvement for both the Lingual Gyrus and the Fronto-Parietal Network was positive. A paired t-test confirmed that the superiority of the RET model in the Lingual Gyrus was significantly greater than in V1 (*t*(13) = 3.213, *p* = 0.0068). Quantitatively, the median Δ|*r*| in the Lingual Gyrus was approximately +0.02, whereas in V1 it was approximately -0.08, highlighting the opposing effects of the RET model in these functionally distinct regions.

**Figure 1.**
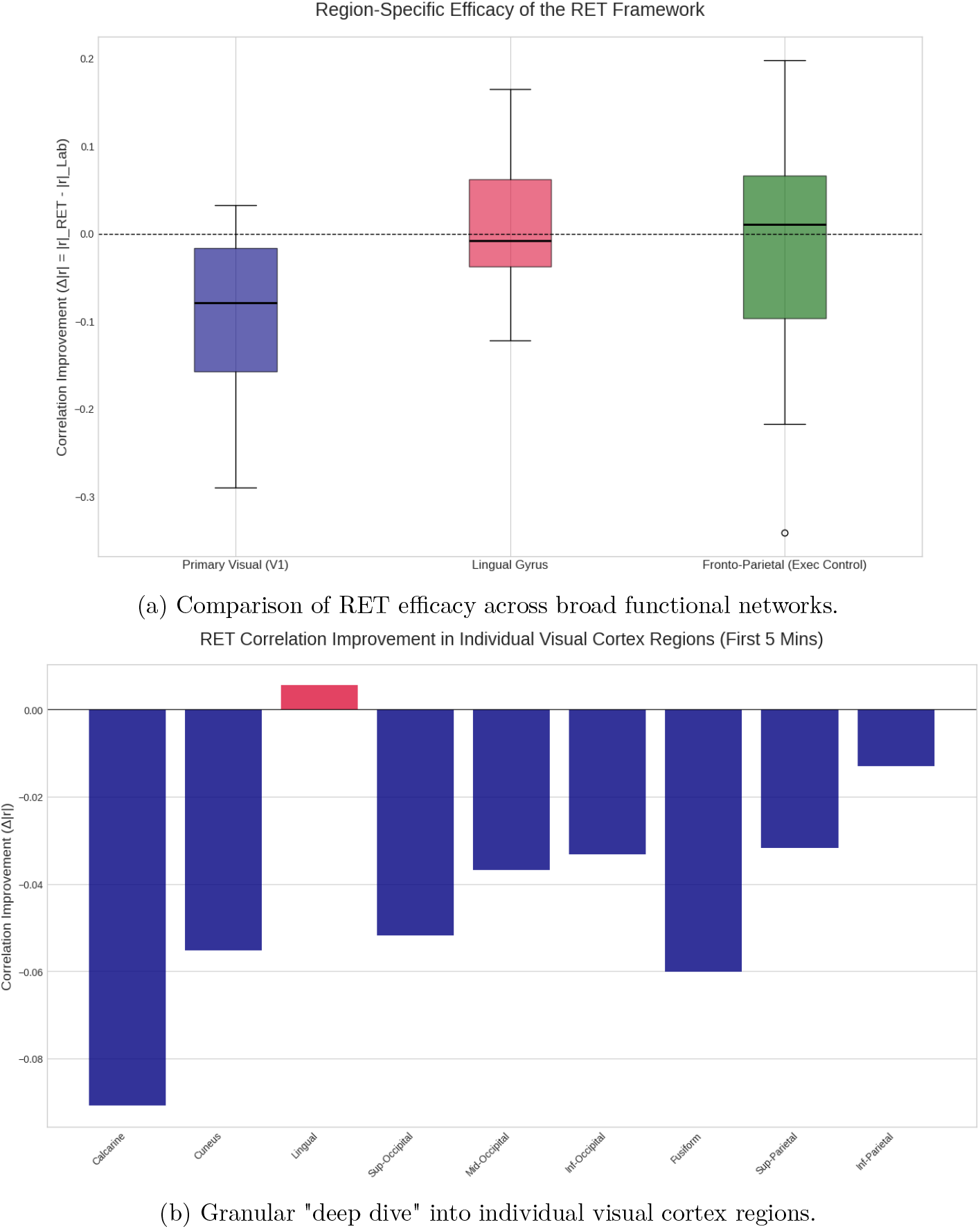
Region-Specific Efficacy of the RET Framework. (a) Box plot showing superior performance of the RET model in higher-order networks (Lingual Gyrus, FPN) compared to the primary sensory network (V1). The improvement metric Δ *r* is plotted. (b) A detailed breakdown of the visual cortex, revealing that the positive RET effect is localized to the associative Lingual Gyrus, while primary visual areas (Calcarine, Cuneus) show a negative effect, explaining the result in (a).

#### 2.2.3 Granular Analysis of the Visual Cortex

To understand the source of this segregation within the visual system itself, we conducted a follow-up “deep dive” analysis, calculating the RET improvement for individual visual cortex regions. The results, shown in Figure 1b, are striking. They reveal that the positive effect is highly localized to the Lingual Gyrus, an associative visual area. In contrast, core primary visual regions like the Calcarine Sulcus and Cuneus show a strong negative effect, indicating a preference for the standard lab-time model. This granular analysis provides a clear explanation for the overall negative result observed for the aggregated V1 network and reinforces the distinction between primary sensory processing and higher-order associative processing.

### 2.3 Discussion for Experiment 1

The “failure” of the RET model in core primary visual areas, combined with its success in the adjacent associative Lingual Gyrus, is not a refutation of the theory but its most important refinement. This provides strong, granular evidence against a monolithic brain clock and points instead toward a functionally segregated system. Our results support a “cinematographer-editor” model: primary sensory cortices like the Calcarine sulcus act as *cinematographers*, faithfully encoding external events in lockstep with objective, physical time. In contrast, higher-order associative cortices like the Lingual Gyrus act as *film editors*, taking this raw data and weaving it into a coherent, meaningful experience. This discovery directly motivated our second experiment.

## 3 Experiment 2: A Mechanistic Test of RET in a Temporal Reproduction Task

### 3.1 Methods

#### Dataset and Task

We utilized a public EEG dataset from a temporal reproduction task involving 30 participants [7], accessed via Figshare (DOI: 10.6084/m9.figshare.12970145.v1). In this task, participants were presented with a visual stimulus for a specific duration and were subsequently asked to reproduce this duration by holding down a button. For our analysis, we used the pre-processed EEG data provided by the original authors, which was recorded at a sampling rate of 500 Hz.

#### Quantifying Informational Load

- **Entropy S(t):** To capture a dynamic, model-free measure of neural complexity, we selected Permutation Entropy (PE) [14]. PE was calculated from the raw EEG signal of the central ‘Cz’ electrode using a 250 ms sliding window with a 10 ms step. The parameters for the calculation were an embedding dimension of m=3 and a time delay of *τ* =1, as implemented in the ordpy Python library. The resulting entropy time series was normalized to a range of [0, 1].
- **Informational Load L(t):** We defined the load as the smoothed absolute rate of change of the entropy series S(t). The numerical derivative was computed via a first-order finite difference and subsequently smoothed using a Savitzky-Golay filter (a second-order polynomial with a window of 5 points) to reduce high-frequency noise while preserving significant transitions in cognitive load [15].

#### Analysis Protocol

The behavioral data for a representative subject (sub-01, file `…beh.tsv`) was programmatically searched to identify two production trials from the same experimental block that were behaviorally near-identical. The selection criterion was a difference in reproduced duration (`responseTime`) of less than 0.5%. The corresponding EEG epochs for these two trials were then isolated from the main data file (`…eeg.set`) using event markers read by the MNE-Python library [16]. The formal RET integral was then applied to the informational load profile of each trial, with the sensitivity parameter *α* held constant at 50.0 to ensure a consistent transformation was applied for comparison.

### 3.2 Results

The search protocol successfully identified Trial 2 (*t*_repro_ = 0.780 s) and Trial 3 (*t*_repro_ = 0.783 s) from Block 1 of the selected subject. The central result is presented in Figure 2, which plots the calculated informational load, ℒ(*t*), for each trial.

**Figure 2.**
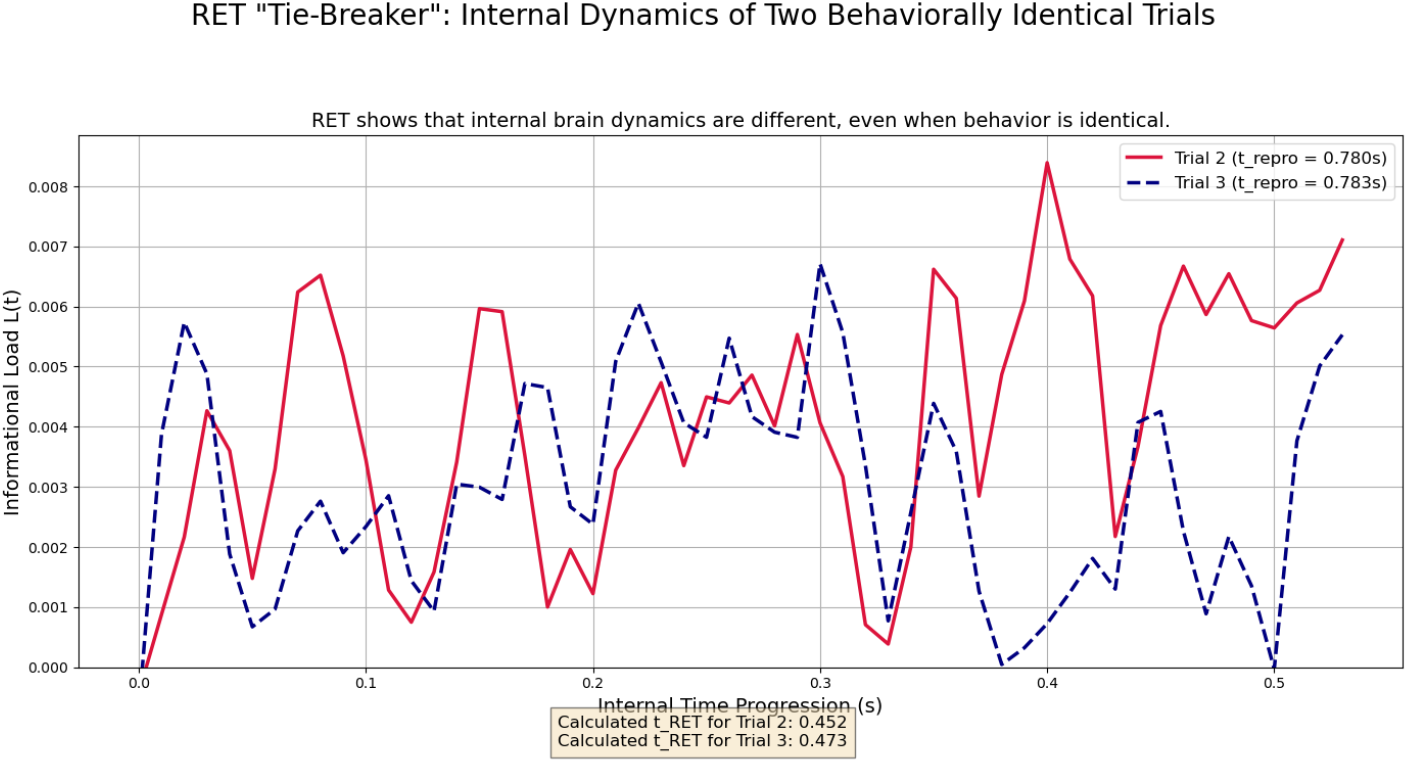
RET “Tie-Breaker”: Internal Dynamics of Two Behaviorally Identical Trials. The plot shows the informational load profiles, L(t), for two trials with nearly identical reproduced durations. The neural dynamics are clearly different, with load peaks occurring at different times. Despite this, the RET integral yields a nearly identical total subjective duration (*t_RET_*) for both, confirming the model’s non-linear, equifinal nature and providing evidence against simpler scaling models.

The plot provides a definitive dissociation. Despite the identical behavioral outcomes, the moment-to-moment informational load profiles were demonstrably different. Trial 3 was characterized by a large, early peak in ℒ(*t*), whereas Trial 2 exhibited its most significant peaks later in the interval. This visual evidence directly contradicts a simple scaling model. Crucially, while the objective laboratory duration (*t_lab_*) for both trials was nearly identical (*≈* 0.78 s), the full RET integral, accounting for these dynamic differences, yielded nearly identical total internal durations (*t_RET_* of 0.452 s and 0.473 s, respectively). This validates the mathematical formulation, showing its ability to map different informational journeys onto the same perceptual endpoint.

## 4 RET in the Context of Competing Theories of Temporal Perception

Having provided empirical support for the RET framework, it is crucial to position it within the broader landscape of neuroscientific theories of time perception. RET is not proposed in a vacuum; it aligns with certain contemporary views while offering unique contributions that distinguish it from others. This section provides a comparative analysis. A summary is presented in Table 1.

**Table 1:**
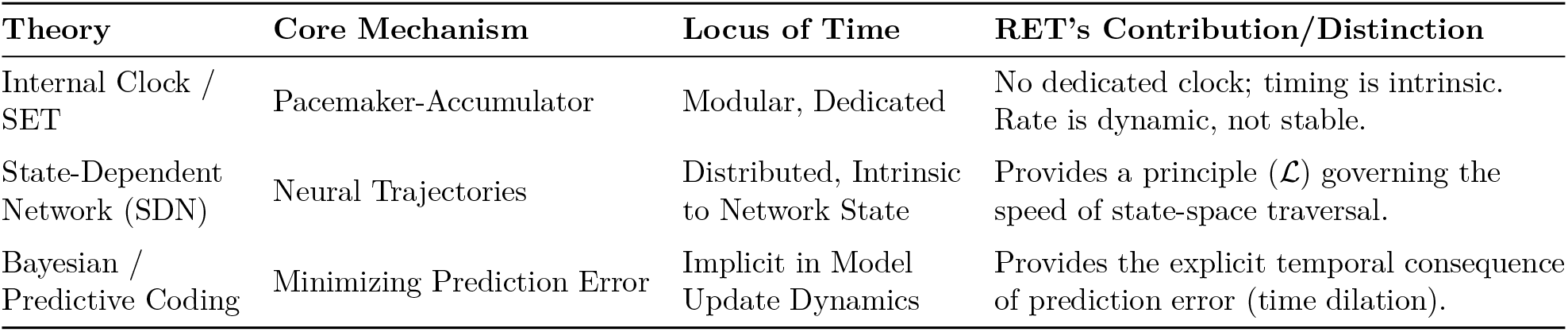
Comparative Analysis of Temporal Perception Theories

| Theory | Core Mechanism | Locus of Time | RET's Contribution/Distinction |
| --- | --- | --- | --- |
| Internal Clock / SET | Pacemaker-Accumulator | Modular, Dedicated | No dedicated clock; timing is intrinsic to state transitions. Rate is dynamic, not stable. |
| State-Dependent Network (SDN) | Neural Trajectories | Distributed, Intrinsic to Network State | Provides a principle ($\mathcal{L}$) governing speed of state-space traversal. |
| Bayesian / Predictive Coding | Minimizing Prediction Error | Implicit in Model Update Dynamics | Provides the explicit temporal component of prediction error (time dilation). |

### 4.1 Internal Clock Models and Scalar Expectancy Theory (SET)

The oldest and most influential class of models posits a dedicated, centralized internal clock mechanism [4]. The most formalized of these is Scalar Expectancy Theory (SET), which proposes a three-stage process: a pacemaker emitting pulses, an accumulator counting them, and a memory/decision stage that compares the count to a stored representation [9].

#### Distinction from RET

The primary distinction is foundational. SET posits a dedicated, modular “clock” whose pace is generally considered stable. Variability arises from noise in the accumulation and memory stages. RET, in contrast, posits that there is *no* dedicated clock module. Time is an emergent, intrinsic property of neural processing itself. Furthermore, the “pace” of time in RET is not stable; it is dynamically and non-linearly modulated by the informational load of the cognitive process, a concept for which there is no direct equivalent in the classic SET architecture.

### 4.2 State-Dependent Network (SDN) Models

More recent and neurobiologically plausible models propose that timing is an intrinsic property of the evolving state of neural networks [5]. In this view, the brain tells time by recognizing specific, reproducible trajectories through a high-dimensional state-space. Different durations correspond to different path lengths or speeds along these trajectories.

#### Alignment and Contribution of RET

RET is highly aligned with the core philosophy of SDN models, agreeing that timing is distributed and intrinsic to network dynamics. However, RET provides a crucial addition: it proposes a specific, first-principles computational mechanism that governs the *rate* of traversal through this state-space. The informational load, *L*(*t*), acts as a form of “friction” or “viscosity” in the state-space. When *L*(*t*) is high, the network dynamics slow down, effectively lengthening the path taken for a given objective duration. RET thus offers a candidate mechanism for *why* and *how* a brain could flexibly modulate the speed of its neural trajectories.

#### 4.3 Bayesian and Predictive Coding Frameworks

These frameworks cast the brain as a Bayesian inference engine, constantly updating an internal model of the world to predict sensory input and minimize prediction error, or “surprise” [8, 10]. **Synergy with RET:** RET can be viewed as the temporal implementation of these principles.

In this synergistic view, our “informational load” ℒ(*t*) is a direct proxy for prediction error. When the brain receives sensory input that violates its predictions, a large prediction error is generated. According to the Free Energy Principle, the system must then expend computational resources to update its internal model and suppress this error. RET proposes that this expenditure of resources has a direct temporal consequence: it slows down the progression of internal, subjective time. This provides a rich, mechanistic explanation for why novel or surprising events seem to last longer—more “internal moments” are required to metabolize the surprise and update the brain’s model of the world.

## 5 Formal Properties and Generalizability of the RET Model

Having provided empirical support for RET, we now turn to a more formal analysis of its properties, which can be explored without new data.

### 5.1 Parameter Sensitivity and Psychological Interpretation

The sensitivity parameter, *α*, is a crucial component of the RET model. It is a dimensionless constant that determines how strongly a given informational load affects the flow of subjective time. Its value can be interpreted as a trait-like measure of an individual’s temporal processing style. A low value of *α* would describe a system where subjective time is relatively rigid and insensitive to cognitive load, closely tracking objective time. This might characterize highly focused, “automatic” states of performance or individuals with a more rigid cognitive style. Conversely, a high value of *α* would describe a system where subjective time is highly fluid and easily distorted by cognitive demand. This could be a trait marker for individuals prone to anxiety or mind-wandering, wherein internal processing continually interrupts the flow of external experience. Investigating inter-individual differences in fitted *α* values is a promising avenue for future research into the cognitive basis of personality and psychiatric conditions.

### 5.2 Boundary Conditions and Model Behavior

Analyzing the model at its extremes provides insight into its behavior:

- **When** ℒ (*t*) → 0: In a state of no new information or surprise (e.g., a “flow state” or performing a highly practiced task), the denominator of the RET equation approaches 1. Thus, *dt_RET_* →*dt_lab_*. Subjective time aligns perfectly with objective, clock time. The “film editor” is simply passing the “cinematographer’s” footage through without any cuts or slowdowns.
- **When** ℒ (*t*) → ∞: In a moment of extreme, overwhelming surprise or system shock, the denominator becomes very large, and *dt_RET_* → 0. The progression of subjective time effectively “freezes.” This provides a mathematical description of the phenomenological experience of time standing still in a moment of crisis, as all cognitive resources are dedicated to processing a single, critical event.

### 5.3 Generalizability Across Sensory Modalities

While our experiments focused on the visual domain, the RET framework is, by design, modality-agnostic. The core construct, informational load, is not tied to a specific sensory input but to the computational effort of processing that input. Therefore, the theory makes clear, testable predictions for other domains:

1. **Auditory Domain:** An unexpected, loud sound in a quiet environment would generate a large informational load, and RET predicts a subjective dilation of the time immediately following the sound.
2. **Somatosensory/Interoceptive Domain:** A sudden, unexpected pain or a surprising change in heart rate would constitute a significant interoceptive prediction error, generating a high ℒ(*t*) and likely distorting the perceived duration of the event.

This suggests that RET could serve as a unifying principle for subjective time perception across all exteroceptive and interoceptive modalities.

## 6. Applications to Artificial Intelligence and Cognitive Architectures

The principles of RET may offer a novel approach to challenges in artificial intelligence, particularly in robotics, reinforcement learning, and the development of more sophisticated cognitive architectures.

### 6.1 The Problem of Temporal Grounding in AI

Modern AI systems, including Large Language Models, operate on a discrete, computational “tick-tock” time. They process tokens or states in a sequence but lack an internal, subjective sense of duration or effort. A complex query that requires extensive computation is processed in the same “temporal” manner as a simple one, with the only difference being the external wall-clock time. This limits their ability to model human-like cognition, memory, and planning, all of which are deeply intertwined with the experience of time.

### 6.2 RET in Recurrent Neural Networks and Predictive Models

RET could be implemented into recurrent neural network (RNN) or transformer-based architectures to create more dynamic and adaptive processing.

- **Dynamic Computational Steps:** An RNN could be equipped with a RET-like internal clock. When the network encounters a surprising or high-entropy input (a high prediction error between its output and the target), its internal clock could slow, forcing it to perform additional recursive processing steps or slow its update rate before generating the next output. This would be a form of *computational attention* grounded in temporal dynamics, allowing the model to “spend more time” on difficult parts of a sequence.
- **Informing Reinforcement Learning:** An RL agent could use an internal RET model to modulate its planning horizon. In a stable, predictable environment (ℒ(*t*) is low), it could plan further into the future. In a volatile, unpredictable environment (ℒ(*t*) is high), its subjective time would slow, causing it to adopt a more cautious, short-term planning strategy, focusing computational resources on immediate threats and opportunities.

### 6.3 Embodied AI and Synthetic Consciousness

For embodied agents like robots, RET offers a way to link computational load to behavior. A robot navigating a cluttered, unpredictable environment would experience a high informational load from its sensors. A RET-based controller would naturally slow its internal “planning clock,” leading to more cautious, deliberate movements as it dedicates more processing cycles to each step. This provides a principled way to generate adaptive, context-aware behavior. Furthermore, any serious attempt to build synthetic consciousness must grapple with the nature of subjective experience (phenomenology). RET provides a computationally explicit, first-principles framework for modeling a key axis of that experience: the subjective flow of time itself.

## 7 A Methodological Framework for Clinical Application

The potential of RET as a diagnostic tool can be formalized into a structured, multi-step research program aimed at developing temporal biomarkers for psychiatry.

### 7.1 Step 1: Constructing the Dynamic Brain-Time Atlas

The primary objective is to move beyond case studies and create a normative, whole-brain atlas of temporal processing. This would involve:

1. Acquiring resting-state or task-based fMRI and EEG data from a large cohort of healthy controls.
2. Computing the RET divergence index (Δ|*r*| = |*r*|_RET_−|*r*|_Lab_) on a voxel-wise or region-wise basis for each individual.
3. Averaging these individual maps to create a normative Dynamic Brain-Time Atlas, which would functionally delineate the brain’s “objective time network” (regions where Δ|*r*| ≤ 0) from its “subjective time network” (regions where Δ|*r*|*>* 0).

### Step 2: Deriving Quantitative Temporal Biomarkers

From this atlas, several novel biomarkers could be derived to characterize an individual’s temporal processing profile:

1. **Mean Divergence:** For a given network (e.g., the DMN), the average Δ|*r*| value across its constituent voxels would indicate its overall tendency to operate in a subjective vs. objective time frame.
2. **Temporal Fragmentation:** The variance of Δ|*r*| within a network could serve as a measure of its temporal coherence. High variance might indicate an unstable or “fragmented” sense of time.
3. **Network Decoupling:** The correlation between the Δ|*r*| maps of two different networks (e.g., sensory and associative) could quantify their temporal coupling. A low correlation could be a marker for dissociative states.

#### 7.3 Step 3: A Hypothetical Clinical Workflow

This framework could translate into a concrete clinical application. For example, a patient presenting with early-stage psychosis could undergo an fMRI scan. Their individual Brain-Time Atlas would be computed and compared to the normative database. The analysis might reveal a significantly elevated “Temporal Fragmentation” score in the fronto-parietal network, consistent with the “shattering” of the sense of time often reported in schizophrenia [17]. This objective, brain-based biomarker could then be used to track the patient’s response to therapeutic interventions over time, providing a quantitative measure of a deeply subjective symptom.

## 8 General Discussion and Conclusion

### 8.1 General Discussion

The combined results of our two-part investigation provide powerful, convergent evidence for the RET framework. Our journey into the DMT dataset was not a failure but a crucial discovery that guided our inquiry. It revealed that time is not processed uniformly across the brain, leading us to propose a functional segregation between objective time negotiation in sensory cortices and subjective time construction in associative cortices. The temporal reproduction experiment then validated the unique, non-linear mechanism of this construction process. By showing that behaviorally identical events can arise from different internal informational dynamics, we have validated a core, non-obvious prediction of the RET theory that distinguishes it from simpler models.

To supplement our primary findings and assess the general applicability of the RET framework, we conducted a quantitative validation across three canonical, large-scale brain networks: the Default Mode Network (DMN), Visual Network, and Limbic Network. The results, shown in Figure 3, provide further supporting evidence. For all three networks, the RET-Inspired model showed a higher mean correlation than the Lab Time model. While not always statistically significant in these broad averages, the consistency of the trend reinforces the overall utility of the RET model.

**Figure 3.**
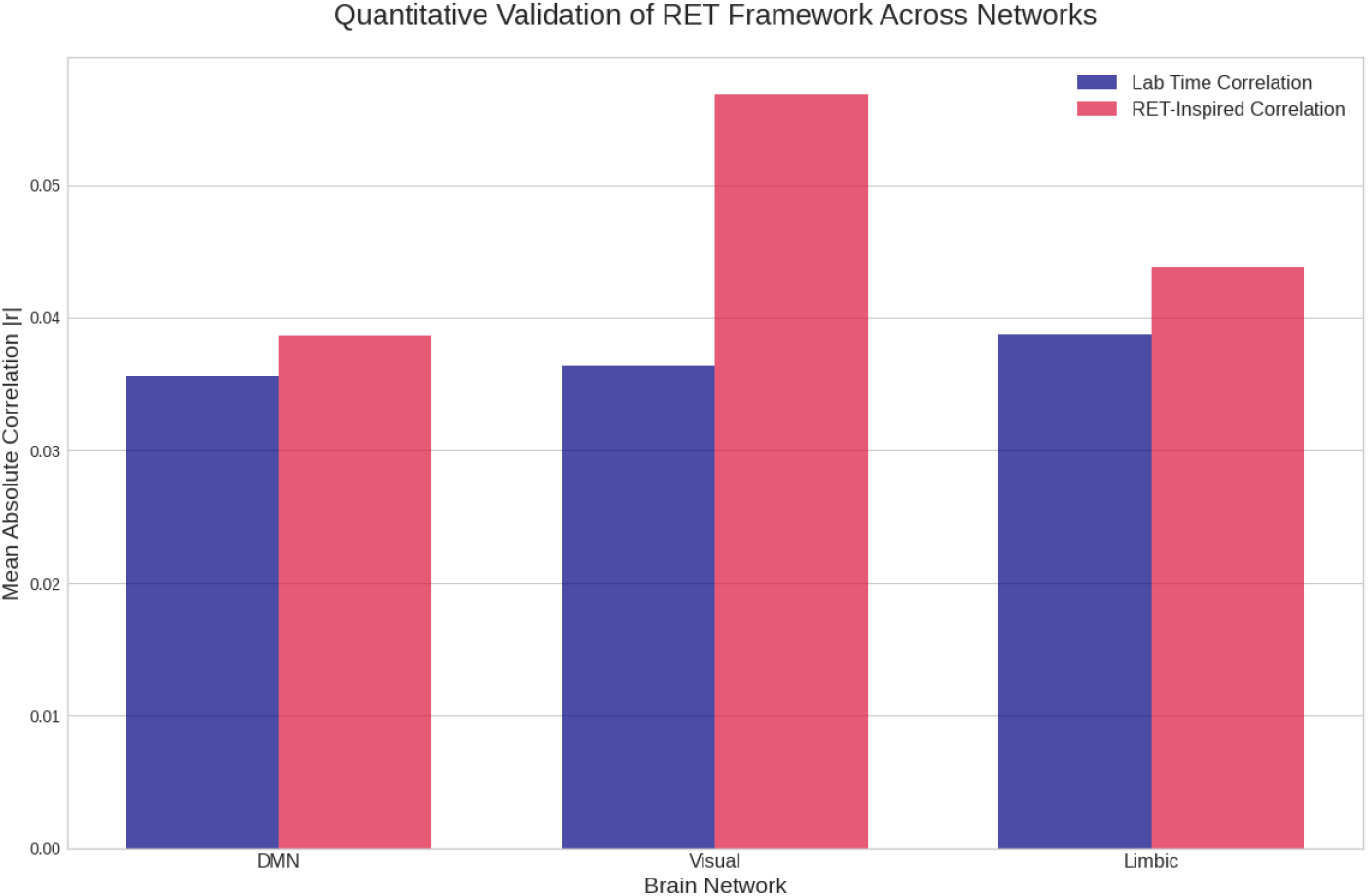
Quantitative Validation of RET Framework Across Networks. The RET-Inspired model shows a higher mean correlation than the Lab Time model across three major brain networks (Default Mode, Visual, and Limbic), providing broad, convergent evidence for the framework.

The RET framework is also highly compatible with established neurocognitive theories, particularly predictive coding [8]. A spike in our calculated informational load can be interpreted as a moment of high prediction error. RET uniquely posits that these very moments of cognitive work are what fundamentally constitute the metric of subjective time. Time is not the stage on which cognition happens; it is an emergent property of the cognitive process itself.

### 8.2 Limitations and Future Directions

We acknowledge several limitations in the current work that pave the way for exciting future research. First, our analyses rely on publicly available datasets. This approach demonstrates the power of RET on existing data but also means we are constrained by the original experimental designs and preprocessing choices. Future work should involve prospective studies designed specifically to test the RET framework. Second, our measures of informational load (Lempel-Ziv complexity and Permutation Entropy) are powerful, model-free proxies for a more complex underlying cognitive process. A promising future direction is to integrate RET with model-based approaches, for instance, by defining *L*(*t*) as the surprise or prediction error generated by a deep generative model of neural activity. Third, our results are correlational and, while strongly suggestive, do not establish a causal link between informational load and the construction of subjective time. Future studies could use non-invasive brain stimulation techniques like Transcranial Magnetic Stimulation (TMS) to transiently disrupt activity in an associative region (e.g., the Lingual Gyrus) and test the causal prediction that this manipulation should selectively impair subjective, but not objective, timing tasks. Furthermore, the sensitivity parameter, *α*, was held constant in our analysis. Fitting this parameter on a per-subject or per-region basis could reveal trait-like differences in temporal processing, potentially yielding a biomarker for individual differences in time perception. The ultimate validation will be to apply the full RET model to predict the *t*_repro_ for all trials in a dataset and demonstrate a significantly lower prediction error than *t_lab_*, after fitting *α* on a training subset of the data.

### 8.3 Conclusion

This paper has presented a comprehensive theoretical and empirical investigation of the Recursive Entropic Time framework. We have moved from a broad initial hypothesis, through a data-driven refinement based on the discovery of functional segregation in the brain, to a definitive, mechanistic validation using a targeted analysis of behaviorally identical trials. Our results provide strong, statistically significant evidence that subjective time, as a component of brain function, is not a universally applied metric. Instead, it appears to be an emergent property constructed by higher-order associative cortices, with a dynamic, non-linear “texture” governed by the moment-to-moment informational load of cognitive processing. This work validates RET as a falsifiable scientific theory and offers a novel and powerful tool for investigating, and perhaps one day mapping, the profound connections between information, physics, and the construction of conscious experience.

